# Uncoupling hypersensitive cell death response and disease resistance activated by effector-triggered immunity

**DOI:** 10.1101/2025.10.22.683861

**Authors:** Himanshu Chhillar, Henk-jan Schoonbeek, Bruno Pok Man Ngou, Jonathan DG Jones, Pingtao Ding

## Abstract

Effector-triggered immunity (ETI) is a major defence strategy in plants and is frequently associated with the hypersensitive response (HR), a localized form of programmed cell death long assumed to be essential for pathogen resistance. However, the causal relationship between HR and effective immunity remains unresolved. We show that the Arabidopsis *cbp60g sard1* double mutant exhibits exaggerated ETI-associated HR but only partial resistance to bacterial and oomycete pathogens, thereby genetically uncoupling cell death from disease resistance without pleiotropic defects. Genome-wide transcriptome profiling reveals that the absence of CBP60g and SARD1 disrupts the balance between immune activators and suppressors, including reduced induction of the Nudix hydrolase NUDT7. Overexpression of *NUDT7* diminishes but does not abolish the heightened HR phenotype in *cbp60g sard1* mutant, indicating that multiple negative regulators act redundantly to restrain immune-associated cell death. These findings demonstrate that HR is not an obligatory determinant of effective resistance and provide mechanistic insight into how plants coordinate transcriptional networks to balance pathogen defence with the containment of host cell death. By refining the relationship between HR and immunity, this work challenges a long-standing paradigm in plant biology and advances our understanding of immune regulation.

## Introduction

Plants are constantly challenged by diverse pathogens that threaten global crop productivity, causing an estimated 20-40% yield loss annually and over US$220 billion in damages according to the Food and Agriculture Organization of the United Nations (FAO) (1). To defend themselves, plants deploy two interconnected layers of innate immunity. Pattern recognition receptors (PRRs) located at the plasma membrane perceive pathogen-associated molecular patterns (PAMPs) to activate pattern-triggered immunity (PTI), while intracellular nucleotide-binding leucine-rich repeat receptors (NLRs) sense pathogen effectors and trigger effector-triggered immunity (ETI) (2). In *Arabidopsis thaliana* (Arabidopsis hereafter), NLRs are classified into coiled-coil (CC)-type (CNLs), Toll/Interleukin-1 receptor/Resistance protein (TIR)-type (TNLs), and RESISTANCE TO POWDERY MILDEW (RPW8)-like CC-type (RNLs) based on their N-terminal domains (3). Although PTI and ETI share many downstream outputs, including calcium (Ca^2+^) influx, reactive oxygen species (ROS) production, and large-scale transcriptional reprogramming, ETI responses are typically more robust and sustained (4).

Calcium (Ca^2+^) signalling is a critical mediator of both PTI and ETI (4). Ca^2+^ signals are sensed by calcium-binding protein calmodulin (CaM) or/and CaM-like (CML) proteins, which interact with downstream CaM-binding proteins (CBPs) to transduce immune signals (5). The Arabidopsis CBP60 family is comprised of eight members, seven of which contain canonical Ca^2+^-dependent CaM-binding domains, whereas SARD1 lacks this domain (6). Among them, CBP60g and SYSTEMIC ACQUIRED RESISTANCE DEFICIENT 1 (SARD1) are pathogen-inducible transcription factors that promote salicylic acid (SA) biosynthesis by directly activating *ISOCHORISMATE SYNTHASE 1* (*ICS1*) and other defence-associated genes (7–10). *cbp60g sard1* double mutants lacking both factors CBP60g and SARD1 display reduced SA accumulation, enhanced susceptibility to the bacterial pathogen *Pseudomonas syringae*, and loss of systemic acquired resistance (SAR) (7, 11). Thus, CBP60g and SARD1 have been recognized as central positive regulators of SA-mediated immunity.

A hallmark of ETI is the hypersensitive response (HR), a localized form of programmed cell death that often accompanies disease resistance (12). HR in plants shared some morphological features with animal cell death (e.g., cytoplasmic shrinkage, chromatin condensation) but also exhibits plant-specific traits such as vacuolization and chloroplast disruption (13). While HR and pathogen resistance are often tightly correlated, several genetic studies indicate they can be uncoupled. For example, loss of the metacaspase in Arabidopsis *mc1* drastically reduces HR but does not abolish resistance (14), and mutants such as *dnd1*, *hlm1*, and *lsd1* maintain resistance despite altered or spontaneous cell death (15–17). However, these mutants often show pleiotropic effects, complicating interpretation (18).

Here, we identify the *cbp60g sard1* double mutant as a valuable model to dissect the relationship between disease resistance and HR. Unlike other uncoupling mutants, *cbp60g sard1* exhibits enhanced HR in the absence of growth defects, providing a genetically tractable system to separate cell death from resistance. We demonstrate that CBP60g and SARD1 contribute to ETI-mediated resistance mediated by different types of NLRs, while paradoxically restraining ETI-associated HR. In addition, we test their contribution to necrotrophic pathogen interactions to gain a broader understanding of its immune function.

By integrating genome-wide transcriptomic profiling with functional assays, we uncover that CBP60g and SARD1 serve as dual regulators of the immune transcriptome: they promote defence-related gene activation while simultaneously maintaining negative regulatory circuits, including members of the Nudix hydrolase family (NUDT). Functional analysis of *NUDT7* demonstrates its role in buffering ETI-associated cell death but also reveals that its overexpression cannot fully rescue the enhanced HR of *cbp60g sard1* double mutant. Together, our findings establish CBP60g and SARD1 as master integrators of positive and negative immune regulation, and provide mechanistic insight into how plants fine-tune immune output, while decoupling HR from disease resistance.

## Materials and Methods

### Plant material and growth conditions

*Arabidopsis thaliana* accessions Col-0 and a β-estradiol (E2) inducible Super-ETI line (SETI_WT), were used as the wild-type controls in this study. Mutants of *eds1-2*, eds1-12, *sard1-1 cbp60g-1* that were used in this study have been previously described (7, 19, 20). Mutants of *eds1-2* and *sard1-1 cbp60g-1* were crossed with SETI_wt to generate SETI_*eds1-2* and SETI_*gh* plants respectively. NUDT7^OE^ overexpression plants in different genetic backgrounds were generated by generating and transforming *AtRBCS1A::gDNANUDT7*-mEGFP-tHSP18.2 construct into the SETI_WT and SETI_*gh*. Plants were grown at 21°C under long-day conditions (16 h light, 8 h dark), and at 50% humidity.

### Bacterial strains and growth conditions

*Pseudomonas syringae* pv. tomato (*Pst*) DC3000 EV (carrying empty vector) grown on the King’s B medium plates containing 25 μg ml^−1^ rifampicin, and 50 μg ml^−1^ kanamycin and *Pst* DC3000 hrcC^−^ were grown on the King’s B medium plates containing 25 μg ml^−1^ rifampicin. *Pseudomonas fluorescens* engineered with a Type III secretion system (Pf0-1 ‘EtHAn’ strains) expressing empty vector or AvrRpt2 was grown on the King’s B medium plates with 50 μg ml^−1^ kanamycin, 34 μg ml^−1^ Chloramphenicol and 5 μg ml^−1^ tetracycline. *Pseudomonas fluorescens* engineered with a Type III secretion system (Pf0-1 ‘EtHAn’ strains) expressing AvrRps4 was grown on the King’s B medium plates with 10 μg ml^−1^ Gentamycin, 34 μg ml^−1^ Chloramphenicol and 5 μg ml^−1^ tetracycline. All the Pseudomonas strains were grown on plates at 28°C for 2 days for further inoculum preparation.

### Hypersensitive cell death response phenotyping in Arabidopsis

Pseudomonas fluorescens engineered with a Type III secretion system (Pf0-1 ‘EtHAn’ strains) expressing one of the effectors (AvrRps4 or AvrRpt2), or pVSP61 empty vector were grown on selective KB plates for 24 h at 28 °C (21, 22). Bacteria were harvested from the plates, resuspended in infiltration buffer (10 mM MgCl_2_), and the concentration was adjusted to OD600=0.2 (108 CFU ml^−1^). The abaxial surfaces of 5-week-old Arabidopsis leaves were hand infiltrated with a 1 ml needleless syringe. Cell death was monitored 3dpi after infiltration.

### Electrolyte leakage assay

Two leaves of 5-week-old *Arabidopsis* plants were hand infiltrated using a 1-mL needleless syringe with 50 μM E2 dissolved in Milli-Q water or Pf0-1 ‘EtHAn’ strain carrying AvrRps4 or AvrRpt2 dissolved in 10mM Mgcl2 (OD600=0.2). DMSO in Milli-Q water or 10mM Mgcl2 was used as mock treatment. Leaf discs were collected with a 7-mm diameter cork borer from infiltrated leaves on paper towels. Leaf discs were dried and transferred into 2 mL of deionised water in 12-well plates (2 leaf disks per well. The plate was incubated for 30 min in a growth chamber with controlled conditions at 21°C under long-day conditions (16-h light/8-h dark) with a light intensity of 120–150 μmol m^−2^. The water was replaced after incubation with 2 mL of deionised water. Electrolyte leakage was measured with Pocket Water Quality Meters (LAQUAtwin-EC-33; Horiba) calibrated at 1.41 mS/cm. Around 100 μL of the sample was used to measure conductivity at the indicated time points. ANOVA (*p* ≤ 0.05) was used for identifying significant factors. Tukey-HSD-Test (*p* ≤ 0.05) was used to determine differences between treatment and lines. A detailed statistical summary is available on GitHub: https://github.com/chhilla/HR_gh.

### Bacterial growth assay

*Pseudomonas syringae* pv. tomato (*Pst*) strain DC3000 carrying pVSP61 empty vector or AvrRps4 and *Pst* DC3000 hrcC^-^ was grown on selective King’s B (KB) medium plates. Bacteria were harvested from the plates, resuspended in infiltration buffer (10 mM MgCl2), and the concentration was adjusted to an optical density of 0.001 at 600 nm [OD600=0.001, representing ∼5×10^5^ colony-forming units (CFU) ml–1]. Bacteria were infiltrated into abaxial surfaces of 5-week-old Arabidopsis leaves with a 1 ml needleless syringe. For quantification, leaf samples were harvested with a 6 mm diameter cork borer (Z165220; Merck-Sigma-Aldrich), resulting in leaf discs with an area of 0.283 cm^2^. Two leaf discs per leaf were harvested as a single sample. For each condition, four samples were collected immediately after infiltration as ‘day 0’ samples to ensure no significant difference introduced by unequal infiltrations, and six samples were collected at 3 dpi as ‘day 3’ samples to compare the bacterial growth between different genotypes, conditions, and treatments. For ‘day 0’, samples were ground in 200 μl of infiltration buffer and spotted (10 μl per spot) on selective KB medium agar plates to grow for 48 h at 28 °C. For ‘day 3’, samples were ground in 200 μl of infiltration buffer, serially diluted (5, 50, 500, 5000, and 50 000 times), and spotted (6 μl per spot) on selective KB medium agar plates to grow for 48 h at 28 °C. The number of colonies (CFU per drop) was monitored, and bacterial growth was represented as CFU cm^−2^ of leaf tissue. All results are plotted using ggplot2 in R (23), ANOVA (p ≤ 0.05) was used for identifying significant factors. Tukey-HSD-Test (p ≤ 0.05) was used to determine differences between treatment and lines. A detailed statistical summary is available on GitHub: https://github.com/chhilla/HR_gh.

### Lesion diameter measurement

**(A)** B. cinerea strain B05.10 was grown as described previously (24) and used for inoculations as described by (25). Arabidopsis wild type and mutant plants were droplet inoculated with B. cinerea at 2.5 × 10^5^ spores/ml in 5 μl of 6 g/L potato dextrose broth, and lesion diameter was measured after 72–96 h. ANOVA (p ≤ 0.05) was used for identifying significant factors. Tukey-HSD-Test (p ≤ 0.05) was used to determine differences between treatment and lines. A detailed statistical summary is available on GitHub: https://github.com/chhilla/HR_gh.

### ROS burst assay

Leaf discs (6 mm diameter) were collected from 5-week-old plants using a cork borer and placed abaxial side down in 96-well plates containing 200 μl of deionized water. Plates were incubated overnight in the dark. The following day, 200 μl of a reaction mixture containing 20 mM luminol (Sigma-Aldrich, A8511), 0.02 mg ml⁻¹ horseradish peroxidase (Sigma-Aldrich, P6782), and 100 nM flg22 was added to each well. Reactive oxygen species (ROS) production was quantified using TECAN microplate reader. ANOVA (p ≤ 0.05) was used for identifying significant factors. Tukey-HSD-Test (p ≤ 0.05) was used to determine differences between treatment and lines. A detailed statistical summary is available on GitHub: https://github.com/chhilla/HR_gh.

### Reverse transcription–quantitative PCR (RT–qPCR) for measuring relative gene expression

For gene expression analysis, RNA was isolated from 5-week-old Arabidopsis leaves and used for subsequent RT–qPCR analysis. RNA was extracted with a VeZol-pure total RNA isolation Kit (RC202-01; Vazyme) and treated with RNase-free DNase (4716728001; Merck-Roche). Reverse transcription was carried out using HiScript III all-in-one RT SuperMix (R333-01; Vazyme). qPCR was performed using a CFX96 Touch™ Real-Time PCR Detection System. Primers for qPCR analysis of *NUDT7* are (F:5’-CTTGCAAGCTAAGTGGATGC-3’; R: 5’-GCGATACTTTAAGGCGCTTG-3’) (11). The elongation factor 1α (EF1α) primers (F: 5’-CAGGCTGATTGTGCTGTTCTTA-3’; R: 5’-GTTGTATCCGACCTTCTTCAGG-3’) were used as an internal control. Data were analysed using the double delta Ct method (26). All results are plotted using ggplot2 in R. A detailed statistical summary is available on GitHub: https://github.com/chhilla/HR_gh.

### RNA-seq raw data processing, alignment, quantification of expression, and data visualization

Pseudomonas fluorescens engineered with a Type III secretion system (Pf0-1 ‘EtHAn’ strains) expressing AvrRps4 and AvrRps4KRVY135-138AAAA was infiltrated in 5- to 6-week-old *cbp60g sard1* double mutant *and* wild-type (Col-0) Arabidopsis leaves using a 1 mL needleless syringe. For RNA-seq analysis, two leaves per plant were collected at 4 hours post infiltration (hpi) as one biological replicate. Untreated samples were included as controls. Total RNA was extracted using the Zymo RNA extraction kit.

The RNA sample was sequenced by Novogene. Raw reads were trimmed into 390 bp clean reads by the Novogene bioinformatics service. At least 12 million paired-end clean reads for each sample were provided by Novogene for RNA-seq analysis. All reads passed FastQC before the following analyses. All clean reads were mapped to the to a comprehensive Reference Transcript Dataset for Arabidopsis Quantification of Alternatively Spliced Isoforms (AtRTD2_QUASI) containing 82,190 non-redundant transcripts from 34,212 genes via Galaxy and Salmon tools (27, 28). The estimated gene transcript counts were used for differential gene expression analysis and statistical analysis with the 3D RNA-seq software (29). The low-expressed transcripts were filtered if they did not meet the criteria of ≥3 samples with ≥1 count per million reads. The batch effects between three biological replicates were removed to reduce artificial variance with the RUVSeq method (30). The expression data were normalised across samples with the TMM (weighted trimmed mean of M-values) (31). The significance of expression changes in the contrasting groups ‘Col-0_kv vs. Col-0_un, groups ‘Col-0_a4 vs. Col-0_un’, ‘*gh*_kv vs. *gh*_un’ and ‘*gh*_a4 vs. *gh*_un’ were determined by the limma-voom method (32, 33). A gene was defined as a significant differentially expressed gene (DEG) if it had a Benjamini–Hochberg adjusted P-value <0.01 and log2[fold change (FC)] ≥ 1 (upregulated) or log2[fold change (FC)] ≤ −1 (downregulated). The GO term analysis was analyzed with g:Profiler (34). Enrichment analysis of transcription factor motifs in the promoters of genes of interest was performed using the AME tool from the MEME Suite (35, 36).

## Results

### The *cbp60g sard1* mutant partially compromises ETI-mediated bacterial resistance but does not impair PTI

To assess the role of CBP60g and SARD1 in effector-triggered immunity (ETI), we challenged the *cbp60g sard1* (*gh* thereafter) double mutant with *Pseudomonas syringae* pv. *tomato* DC3000 strains carrying either AvrRps4 or AvrRpt2, alongside the corresponding empty vector (EV) controls. In wild-type (WT) Co-0, both effectors conferred strong resistance compared with EV infections, whereas *gh* plants showed only partial restriction of bacterial growth (Figures 1A, S1A), in consistence with previous studies (7, 8). AvrRps4 is recognized by the Toll/Interleukin-1 receptor/Resistance protein (TIR)-type nucleotide-binding leucine-rich-repeat receptors (NLRs) (or TNLs) RESISTANCE TO RASTONIA SOLANACEARUM 1 (RRS1)/RESISTANCE TO PSEUDOMONAS SYRINGAE 4 (RPS4) and RRS1B/RPS4B (37, 38), and AvrRpt2 is recognized by the Coiled-coil (CC)-type NLR (or CNL) RPS2 (39, 40). Therefore, the observed reduction of immunity in *gh* was comparable across TNL- and CNL-triggered pathways, indicating CBP60g and SARD1 contribute broadly to ETI-mediated defence. However, resistance was not completely abolished, suggesting residual immune signalling through salicylic acid (SA), N-hydroxy pipecolic acid (NHP), or additional regulatory modules (41).

**Figure 1.**
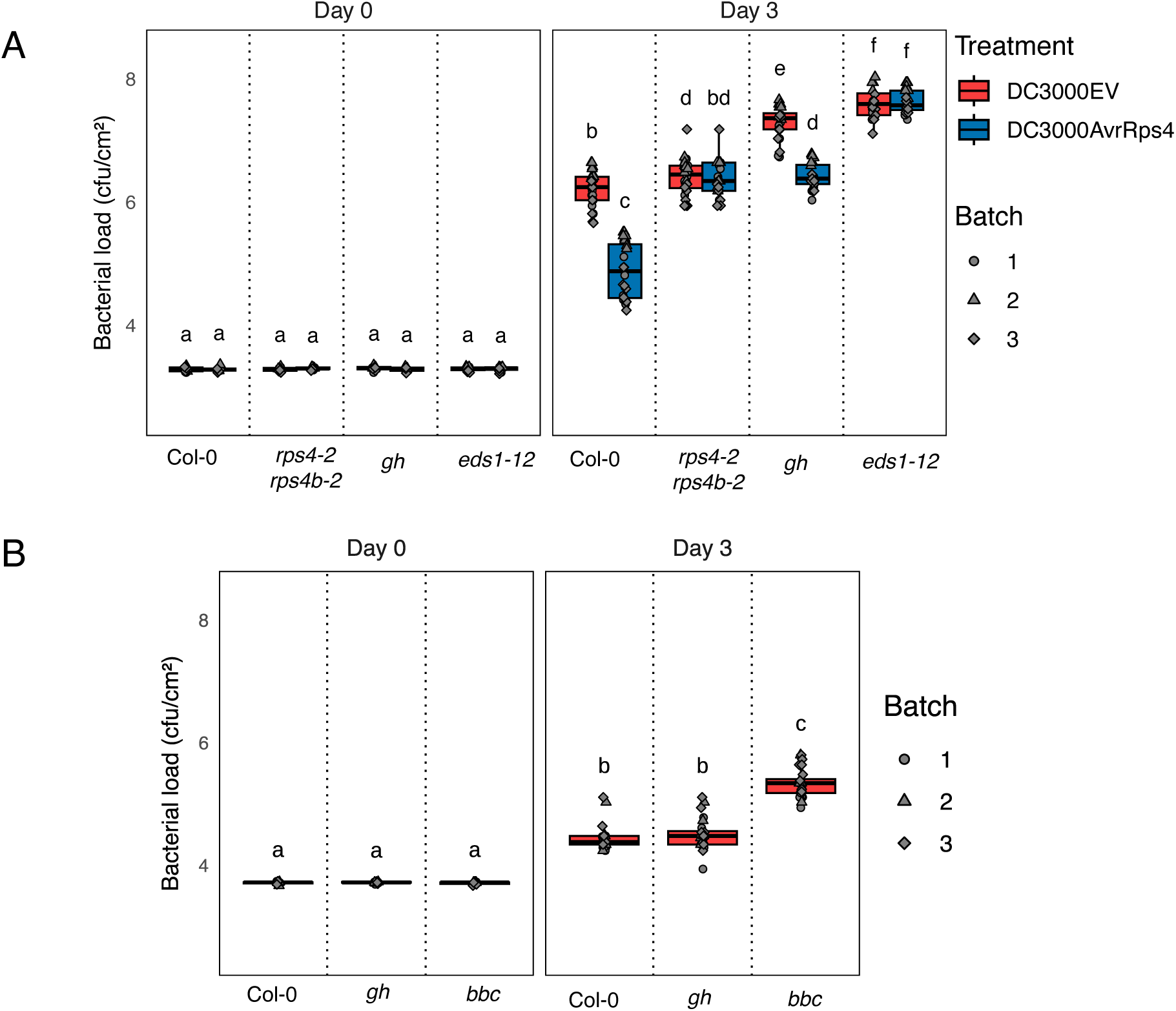
Contribution of CBP60g and SARD1 in PTI and ETI. **(A)** Leaves of Col-0 and the indicated mutant genotypes were infiltrated with *Pseudomonas syringae* pv. *tomato* DC3000 carrying either an empty vector (EV) or AvrRps4. Bacterial growth was quantified as colony-forming units (CFU) at 0 and 3 days post-inoculation (dpi). Statistical differences are indicated by different letters (Tukey’s honestly significant difference [HSD] test, p ≤ 0.05). **(B)** Col-0, *gh* (*cbp60g sard1*), and *bbc* (*bak1-5, bkk1-1, and cerk1*). plants were infiltrated with DC3000 hrcC^-^, and bacterial titres were measured at 0 and 3 dpi. Statistical differences are denoted by different letters (Tukey’s HSD test, p ≤ 0.05).

In contrast, when infected with the non-pathogenic strain DC3000 *hrcC*^-^, which activated pattern-triggered immunity (PTI) but lacks a type-III secretion system, *gh* mutant supported bacterial growth at levels similar to WT Col-0 (Figure 1B). Consistently, *gh* plants displayed WT-like flg22-induced reactive oxygen species (ROS) bursts (Figures S1B, S1C). These results indicate that the absence of CBP60g and SARD1 are not sufficient to compromise canonical PTI signalling outputs but instead play a more specific and central role in ETI.

Together, these findings establish that CBP60g and SARD1 are broadly required for full ETI-mediated resistance across distinct NLR classes, while their absence leaves PTI early signalling intact. The partial loss of ETI in *gh* mutants highlights both the robustness of immune signalling and the presence of compensatory pathways that operate in parallel to CBP60g and SARD1.

### The *sard1 cbp60g* mutant enhances ETI-induced hypersensitive cell death

The hypersensitive response (HR) is a hallmark of effector-triggered immunity (ETI), so we examined whether CBP60g and SARD1 contribute to the regulation of ETI-associated cell death. When infiltrated with *P. fluorescence* Pf0-1 ‘EtHAn’ strains expressing AvrRps4 or AvrRpt2 (21, 42), *gh* mutants showed markedly stronger macroscopic HR symptoms compared with WT Col-0 (Figure 2A). This phenotype was corroborated by electrolyte leakage assays, which revealed significantly elevated ion leakage in *gh* relative to WT following both TNL- and CNL-mediated (AvrRps4 and AvrRpt2, respectively) recognition (Figures 2B-D, S2A, S2B). Enhanced HR in *gh* was evident as early as 22 hours post infiltration (hpi) and was consistently stronger than in Col-0 or other control genotypes. These results indicate that CBP60g and SARD1 normally act to limit the amplitude of ETI-associated cell death across distinct NLR classes.

**Figure 2.**
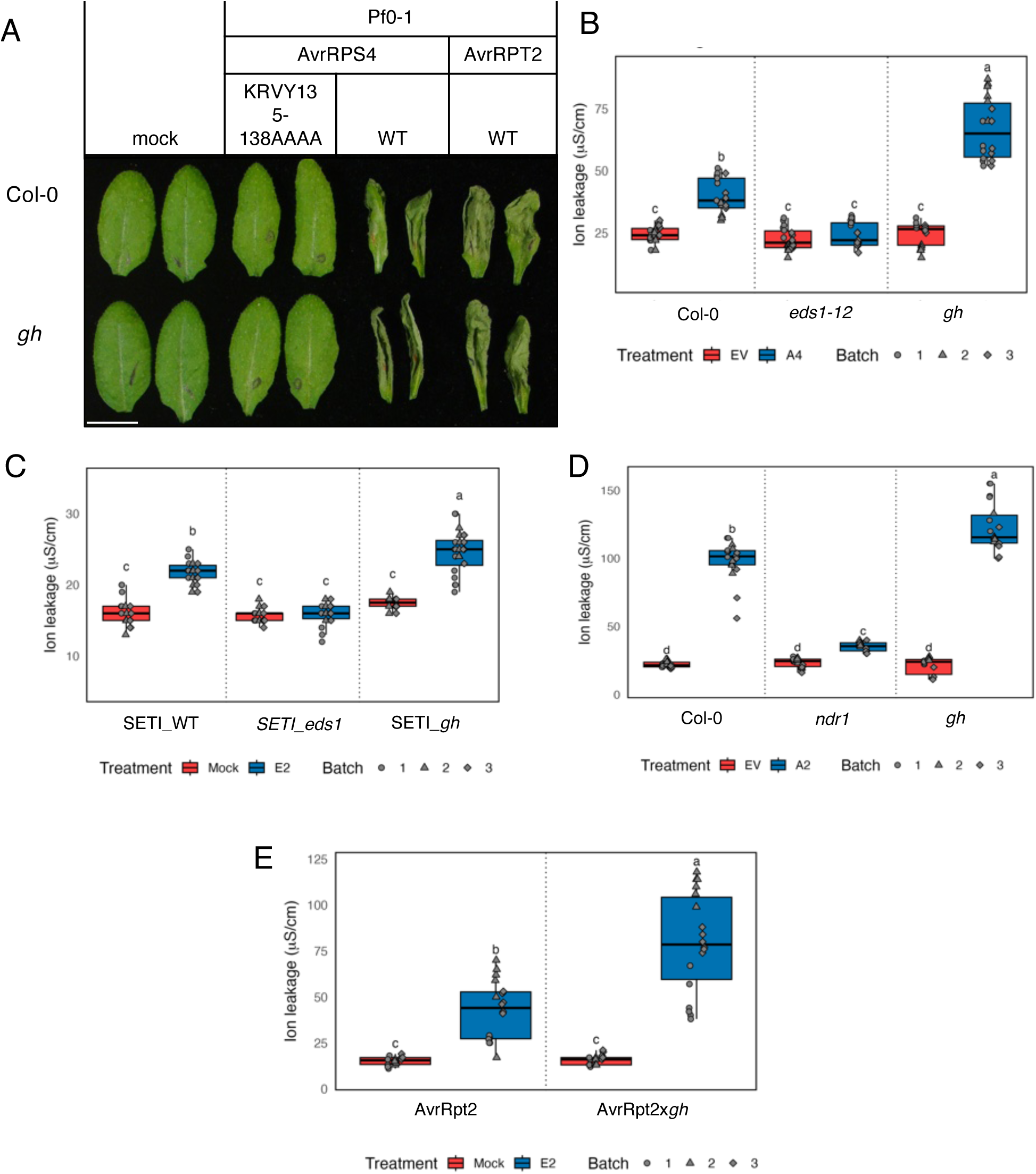
Role of CBP60g and SARD1 in ETI-enhanced HR. **(A)** Leaves of the indicated genotypes were infiltrated with *Pfo-1* strains expressing AvrRps4 ^WT^, AvrRps4 ^KRVY135–138AAAA^, or AvrRpt2 and imaged at 1 day post-infiltration (dpi). Scale bar = 1 cm; arrows indicate PopP2 recognition in Col-0 lines overexpressing RRS1-R and RPS4. **(B, C)** Ion leakage assays in the indicated genotypes infiltrated with *Pfo-1* carrying an empty vector or AvrRps4 (B) and with mock or E2 treatment (C). Measurements were taken 3dpi post-infiltration. **(D, E)** Ion leakage assays in the indicated genotypes infiltrated with s*Pfo-1* carrying empty vector or AvrRpt2 (D) and with mock or E2 (E). Measurements were taken 24 h post-infiltration. Different letters indicate statistically significant differences (Tukey’s HSD test, p ≤ 0.05).

To exclude potential contributions of pattern-triggered immunity (PTI), we analysed HR phenotype in inducible ETI systems (43). When crossed into estradiol-inducible AvrRps4 (the Super ETI or SETI line) or AvrRpt2 plants (43, 44), *gh* mutants exhibited enhanced cell upon ETI induction but not under mock conditions (Figures 2C-E, S3A, S3B). Moreover, when PTI and ETI were activated separately or in combination (PTI+ETI), SETI_*gh* displayed stronger cell death under both ETI and PTI+ETI compared with SETI_WT, while PTI alone induced no such enhancement (Figure S3C).

Thus, despite reduced pathogen resistance, *gh* plants paradoxically undergo more severe ETI-associated HR. This genetic separation of cell death from disease resistance highlights the distinct regulatory layers governing immune outputs and positions the *gh* mutant as a powerful model for dissecting how plants balance defence activation with the containment of self-damaging responses.

### The *sard1 cbp60g* mutant does not enhance susceptibility to necrotrophic fungi

Necrotrophic pathogens exploit host cell death to promote infection (45, 46). Given the enhanced HR in *gh* mutants, we tested whether this would render plants more vulnerable to the necrotrophic fungus *Botrytis cinerea*. Unexpectedly, *gh* plants displayed lesion sizes comparable to WT Col-0 (Figure 3).

**Figure 3.**
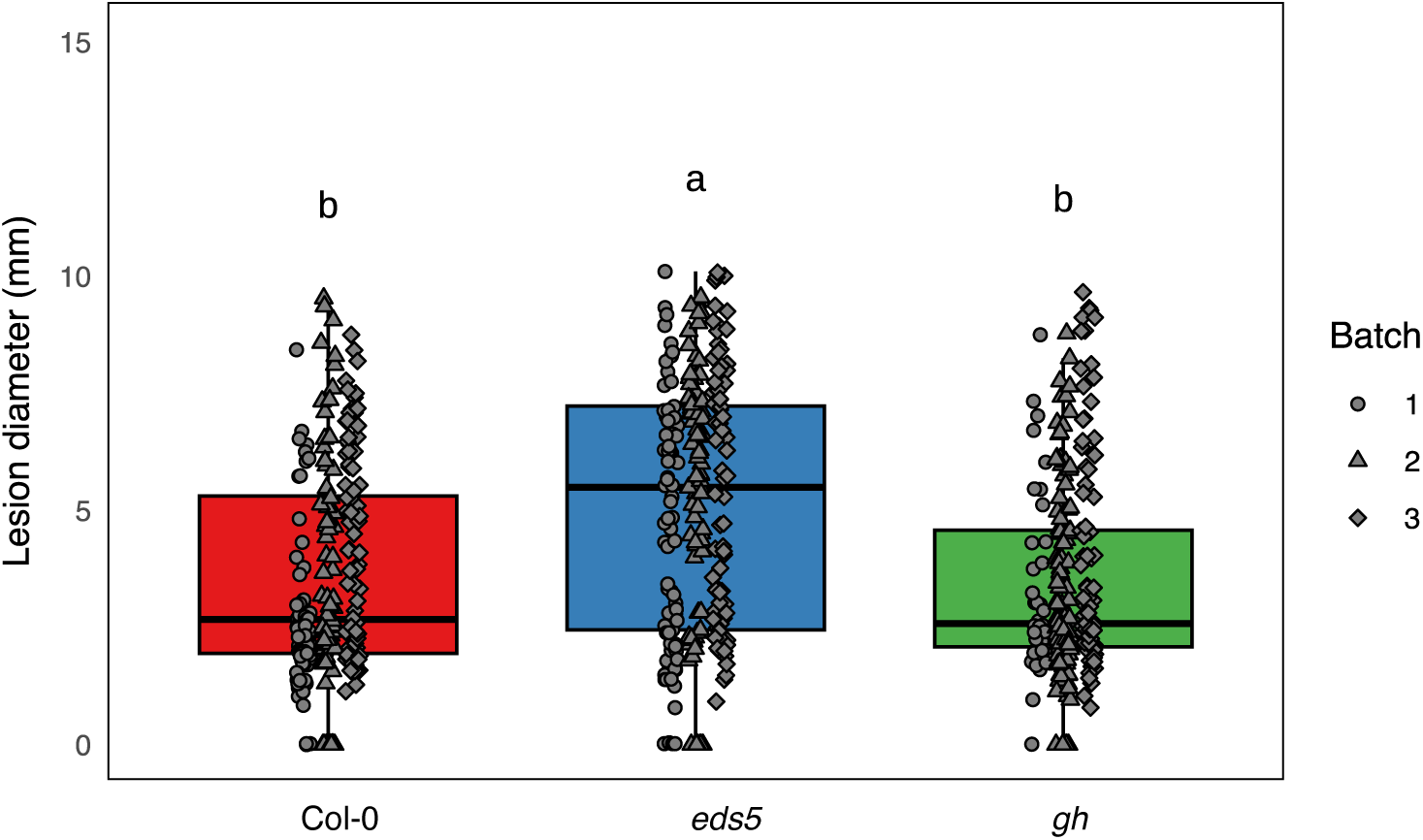
CBP60g and SARD1 do not contribute to resistance against the necrotrophic pathogen *Botrytis cinerea*. The indicated genotypes were droplet-inoculated with B. cinerea strain B05.10, and lesion diameters were measured 72–96 h post-inoculation. Different letters denote statistically significant differences (Tukey’s HSD test, p ≤ 0.05).

In contrast, *eds5* mutants, an established SA-deficient background, showed significantly larger lesions, consistent with their increased susceptibility to *B. cinerea* (Figure 3). Previous work demonstrated that SA-deficient mutants, including *eds5*, undergo enhanced ETI-associated cell death and ion leakage in response to effectors such as AvrRpt2 (47), which correlates with increased vulnerability to necrotrophs. Based on this precedent, we expected *gh* mutants to mirror this phenotype. Instead, *gh* mutants retained WT-like resistance to *B. cinerea* despite their stronger HR.

These findings reveal that enhanced HR in *gh* does not translate into increased susceptibility to necrotrophs, in contrast to SA-deficient backgrounds. This suggests that the cell death program in *gh* is qualitatively distinct, potentially representing a regulated immune-associated pathway that is uncoupled from the necrotrophy-promoted cell death observed in *eds5* and similar mutants.

### The *sard1 cbp60g* mutant disrupts early transcriptional responses to ETI

Early response genes (ERGs) (22, 48) are rapidly induced downstream of both pattern-recognition receptors and NLRs, and many are downregulated in the *gh* double mutant. To dissect the transcriptional programs controlled by CBP60g and SARD1, we performed RNA-seq on Col-0 and *gh* plants challenged with EtHAn:AvrRps4^KRVYmutant^ (PTI) and EtHAn:AvrRps4^WT^ (PTI+ETI) (49).

Across all conditions, we detected 12,174 differentially expressed genes (DEGs, FDR < 0.01, |log₂ fold change| ≥ 1) (Figure 4). Hierarchical clustering grouped these DEGs into 10 major clusters (Figure 4A), highlighting widespread reprogramming of the immune transcriptome upon the loss of CBP60g and SARD1.

**Figure 4.**
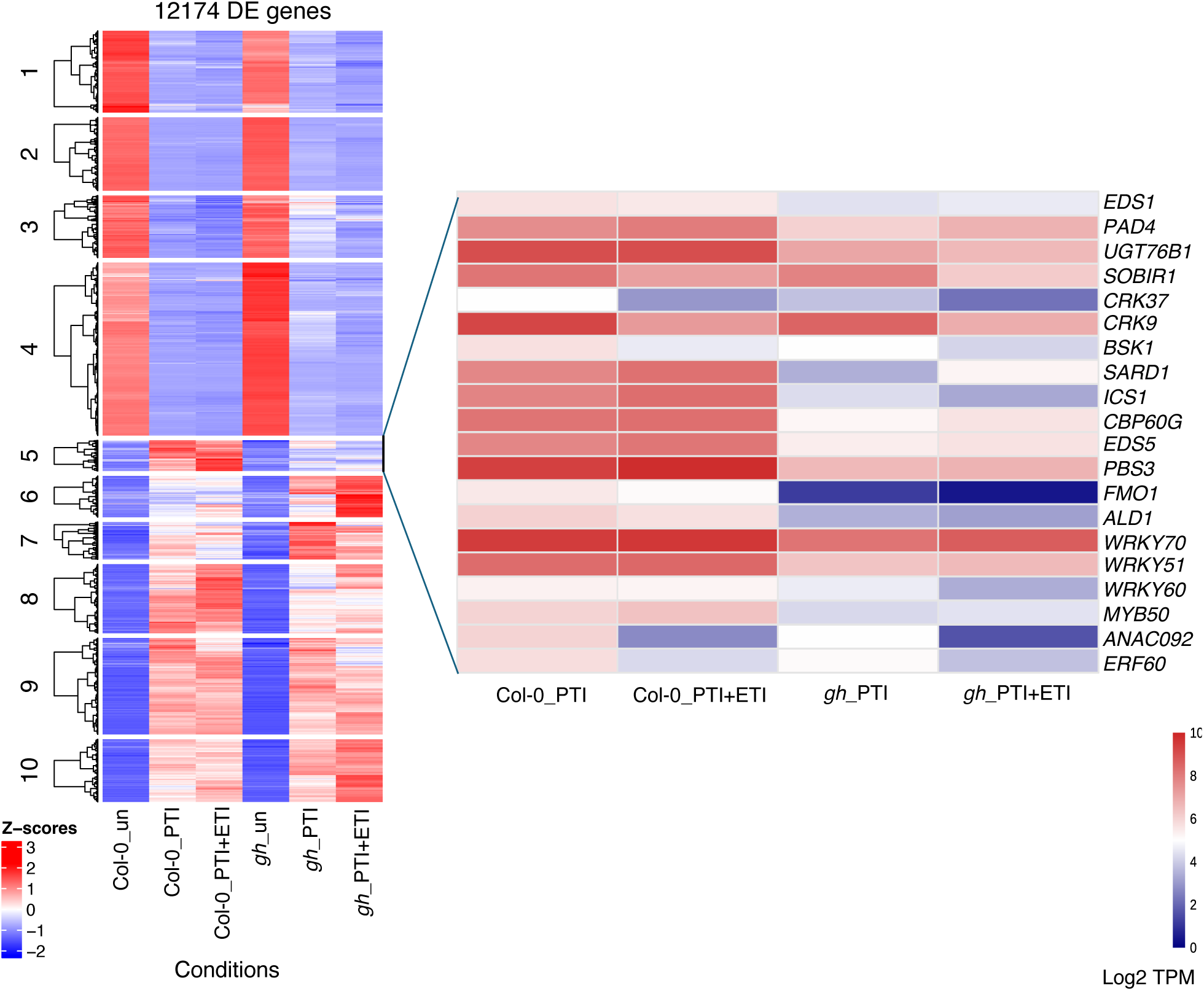
CBP60g/SARD1-dependent transcriptional changes in SETI lipase-like protein mutants. Five-to six-week-old Col-0 and *gh* mutant plants were infiltrated with *Pfo-1* carrying either an empty vector or AvrRps4. Leaf samples were collected 4 h post-inoculation (hpi) for RNA-seq analysis. The left heatmap shows normalized expression (Z-score) of differentially expressed genes (false discovery rate [FDR] < 0.01; log2 fold change ≥ 1). The right heatmap highlights expression patterns of key immune genes from cluster 5 which showed compromised expression in *gh* mutants following immune activation.

Cluster 5 was particularly notable: genes in this group were strongly induced during ETI in Col-0 but showed markedly attenuated expression in *gh* (Figure 4). Gene Ontology (GO) analysis revealed enrichment for defence and metabolic pathways (Figure 5A), including salicylic acid (SA) biosynthesis (*ICS1* and *PBS3*), N-hydroxypipecolic acid (NHP) biosynthesis (*ALD1*, *SARD4* and *FMO1*), the hypersensitive response (HR), and core immune regulators (*EDS1* and *PAD4*). Transcription factors from immunity associated families (WRKYs, ERFs and NACs) were also prominent, indicating a broad defect in immune reprogramming in the *gh* mutant.

**Figure 5.**
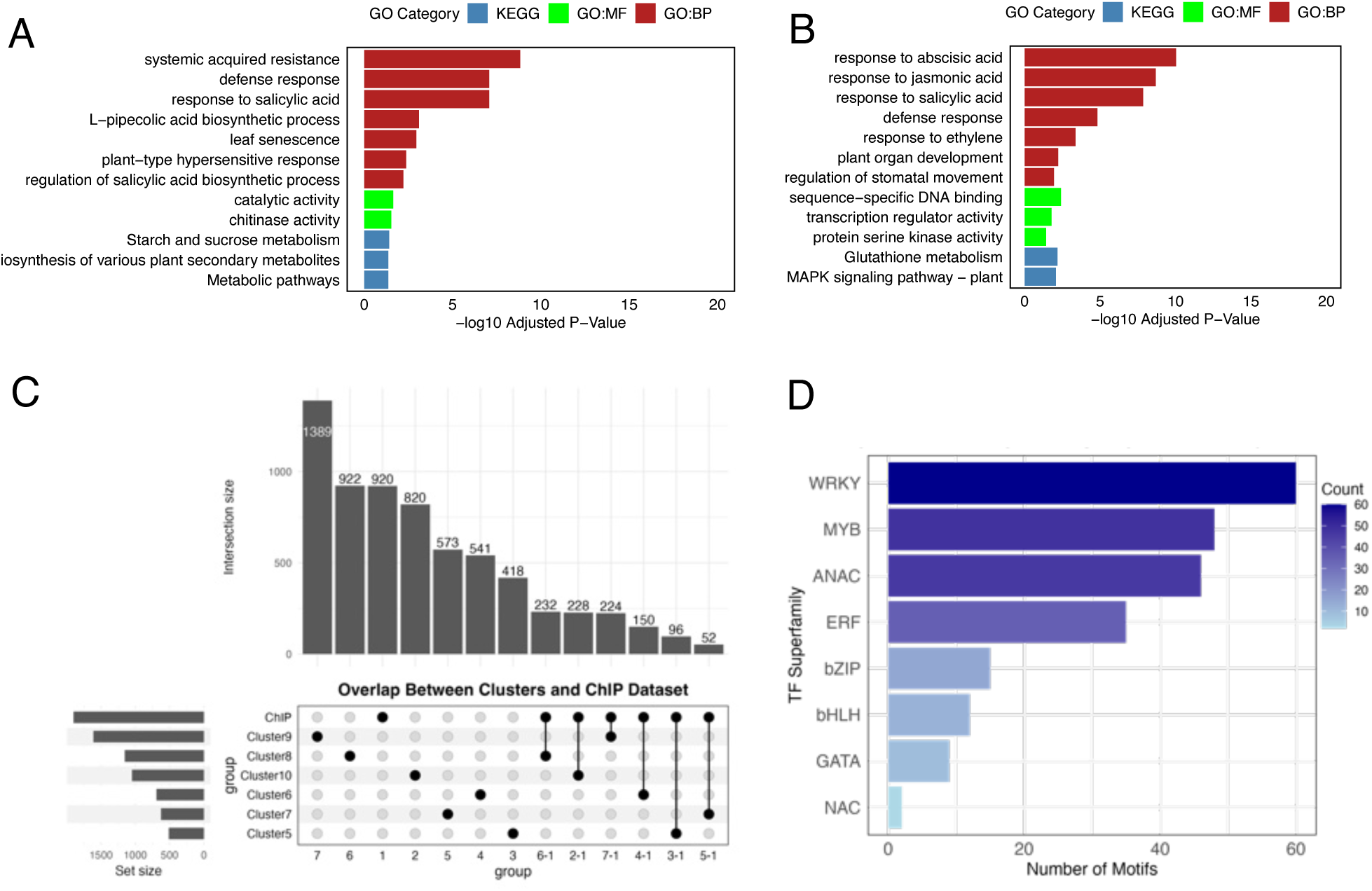
Functional analysis of CBP60g/SARD1-dependent and independent gene clusters. **(A) and (B**) Gene Ontology (GO) enrichment analysis of clusters 5 and 8 was performed using g:Profiler. Enriched GO terms are categorized into biological processes (GO:BP), cellular components (GO:KEGG), and molecular functions (GO:MF) (FDR ≤ 0.05, Benjamini–Hochberg correction). **(B)** UpSet plot showing overlap of differentially expressed genes (DEGs) from various clusters with the SARD1 ChIP-seq dataset reported by Sun et al. (2015). Cluster 9 exhibits minimal overlap with the ChIP-seq dataset. **(C)** Bar graph depicting enrichment of transcription factor motifs in cluster 9 with representation from all major classes such as WRKYs, MYBs, ANACs, ERFs etc.

Cluster 8 showed a similar loss of induction in *gh* and was enriched for hormone- and metabolism-related genes (Figure 5B), including pathways linked to ethylene (ET), jasmonic acid (JA), and abscisic acid (ABA). These data suggest that CBP60g and SARD1 not only control canonical immune regulators but also orchestrate the metabolic adjustments that accompany immune activation.

Together, these findings establish CBP60g and SARD1 as central integrators ETI-induced transcription. Their absence compromises the activation of both immune signalling and metabolic pathways, highlighting their dual role in coordinating transcriptional and physiological reprogramming during plant immunity.

### Dual regulatory roles of CBP60g and SARD1 in the plant immune transcriptome

In addition to their well-established roles as positive regulators of immunity, our transcriptome-scale analysis revealed that CBP60g and SARD1 also exert negative regulatory control over specific gene modules. Hierarchical clustering identified clusters 6 and 10, which contained genes upregulated in the *gh* mutant compared with Col-0, suggesting these genes are normally constrained by CBP60g and SARD1 (Figure S4A, S4B and Table 1). GO enrichment associated these clusters with defence responses, calcium signalling, cell death, and cellular organization, indicating that loss of CBP60g and SARD1 help prevent widespread mis-regulation of physiological and cellular processes.

Importantly, these functions link CBP60g and SARD1 to the control of immune intensity and spatial localization. By repressing signalling modules closely tied to programmed cell death, CBP60g and SARD1 may prevent excessive or ectopic immune activation that could be detrimental to the host. Expression analysis further showed that several well-characterized negative regulators, including members of the Nudix hydrolase family (*NUDTs*), *BAP1*, *BON3*, and *LSD1*, were significantly reduced in *gh* (Figure S4C). While some of these genes were previously identified as CBP60g and SARD1 binding targets (11), our results demonstrate their functional mis-regulation in the mutant background, providing a transcriptome-level view of how CBP60g and SARD1 integrate both positive and negative arms of immune regulation.

Beyond CBP60g- and SARD1-dependent pathways, our data also uncovered modules that appear largely independent of these factors. Cluster 9, for instance, was only modestly affected in *gh* (Figure 4) and showed minimal overlap with the SARD1 ChIP-seq dataset (11), in contrast to the strong overlap observed for clusters 5 and 8 (Figure 5C and Table 2). Motif enrichment of cluster 9 highlighted families such as WRKY, MYB, ANAC, ERF, bZIP, and bHLH, with WRKY motifs being the most enriched (Figure 5D and Table 3). These findings indicate that additional transcription factor networks operate in parallel to CBP60g and SARD1 to drive ETI-associated transcriptional reprogramming.

Together, these results support a model in which CBP60g and SARD1 function as master integrators of the immune transcriptome, coordinating both activation and repression of defence programs. At the same time, the persistence of CBP60g and SARD1independent modules underscores a layered regulatory architecture that balances robust defence activation with the containment of self-damaging responses.

### Overexpressing of *NUDT7* only partially suppresses enhanced ETI-associated HR in *cbp60g sard1* mutants

We hypothesized that the enhanced HR in *gh* plants arises from the misregulation of transcriptional repressors, leading to an imbalance between immune activation and suppression. Previous studies suggested that CBP60g and SARD1 finetune ETI in part by promoting the expression of negative regulators, including members of the Nudix hydrolase family (50). Nudix hydrolases hydrolyse nucleoside diphosphates linked to other moieties (X), thereby maintaining nucleotide homeostasis and modulating stress responses (51). This enzymatic activity is conserved across all domains of life, with 29 homologs identified in *Arabidopsis thaliana* (Figure S4D), underscoring their central role in cellular regulation.

To explore their contribution to ETI regulation, we profiled Nudix hydrolase family members in responses to EtHAn:AvrRps4 (Figure S4D). Among them, *NUDT7* was the most strongly induced in Col-0 but showed markedly reduced induction in *gh* (Figure 6A), indicating that CBP60g and SARD1 are required for ETI-dependent transcriptional activation of this key negative regulator. Consistent with its proposed function, *NUDT7* has been shown to suppress TIR-only protein RESPONSE TO THE BACTERIAL TYPE III EFFECTOR PROTEIN HOPBA1 (RBA1)-mediated cell death in *Nicotiana benthamiana* (52), supporting its conserved role as a negative regulator of immune-associated HR.

**Figure 6.**
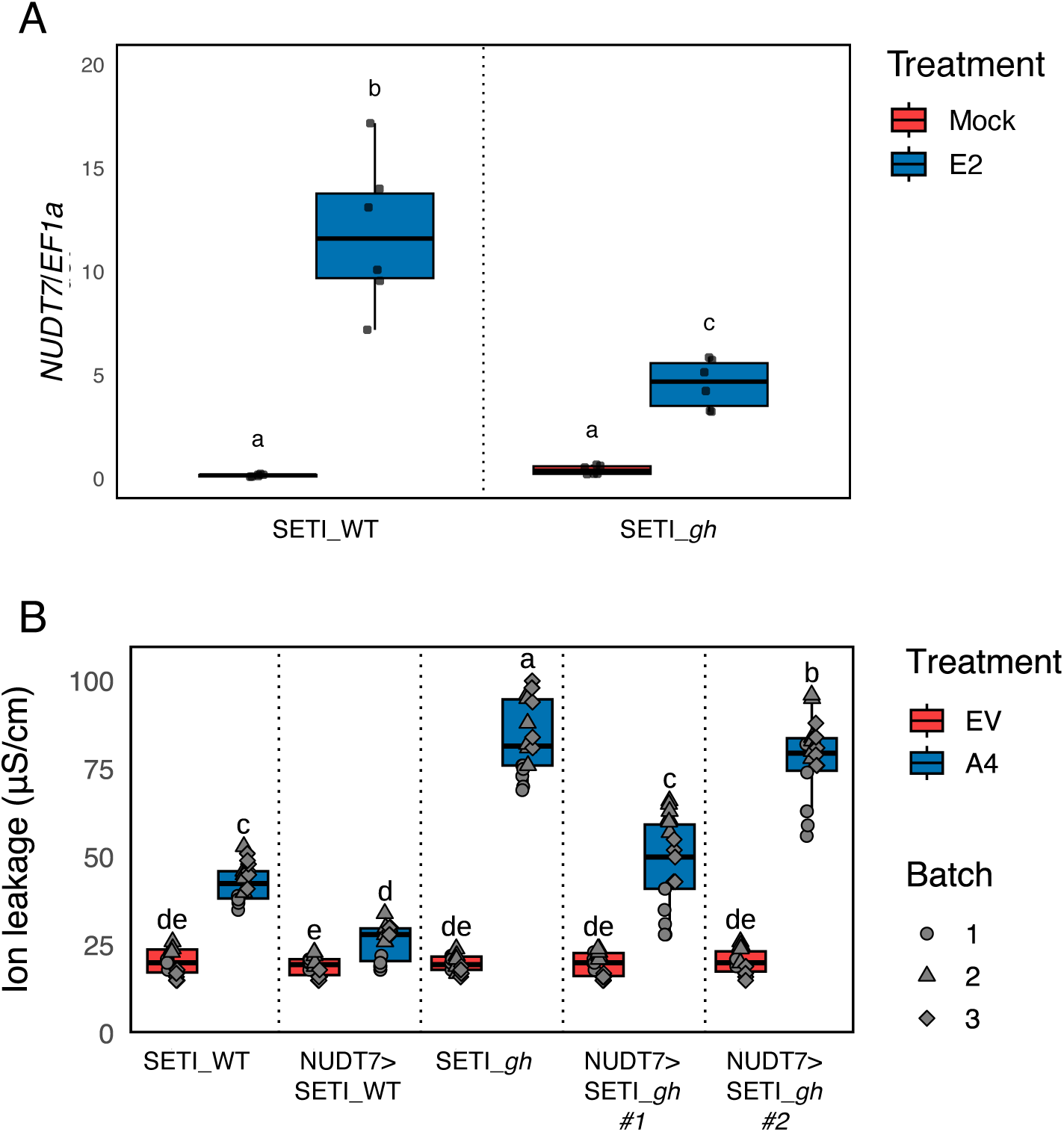
Contribution of NUDT7 to ETI-induced cell death. **(A)** Five-week-old SETI_WT and SETI_*gh* plants were infiltrated with mock or 50 μM E2. Samples were collected 4 h post-infiltration (hpi) for RNA extraction and qPCR analysis. *NUDT7* expression levels are shown relative to *EF1α*. Different letters indicate significant differences (Tukey’s HSD test, p ≤ 0.05). **(B)** Ion leakage assays of the indicated genotypes infiltrated with *Pfo-1* carrying either an empty vector or AvrRps4. Measurements were taken 24 hpi. Statistical differences are denoted by different letters (Tukey’s HSD test, p ≤ 0.05).

To test whether restoring *NUDT7* expression could compensate for the loss of CBP60g and SARD1, we generated *NUDT7* overexpression (*NUDT7*^OE^) lines in both SETI_WT and SETI_*gh* backgrounds (Figure S5A). Upon infiltration with EtHAn_AvrRps4, *NUDT7*^OE^ plants exhibited reduced HR compared with their non-transgenic counterparts, confirming a suppressive role for *NUDT7* (Figures 6B, S5B and S5C). However, in the *gh* background the enhanced HR was only partially suppressed, indicating that NUDT7 overexpression alone cannot restore immune homeostasis that is caused by the loss of CBP60g and SARD1.

These results demonstrate that although NUDT7 contributes to buffering ETI-associated cell death, the severe HR phenotype in *gh* mutants reflects broader misregulation of multiple negative immune regulators beyond NUDT7. Thus, CBP60g and SARD1 act at the network level to balance immune activation with transcriptional repression, and loss of this dual control cannot be compensated by a single negative regulator.

## Discussion

CBP60g and SARD1 are widely recognized as transcriptional activators of plant immunity, acting upstream of SA biosynthesis and systemic signalling. Here, we show that their contribution extends across both TNL- and CNL-mediated ETI, as *cbp60g sard1* (gh) double mutants displayed reduced resistance to bacterial carrying AvrRps4 and AvrRpt2. Notably, while SARD1 directly binds to the promoters of *EDS1* and *PAD4*, both of which encode central components of TNL signalling, it does not bind to *NDR1*, encoding the CNL adaptor, and *NDR1* expression is unchanged in *gh* compared to the WT. The observation that both pathways are equally compromised suggests that CBP60g/SARD1 also regulate shared downstream immune modules that act as convergence points between NLR classes, in addition to their established role in SA biosynthesis.

Although CBP60g and SARD1 bind promoters of PTI-associated genes and are transcriptionally induced upon PAMP treatment, *gh* mutants did not exhibit altered resistance against the *hrcC*⁻ mutant, which elicits canonical PTI. These findings argue that CBP60g and SARD1 are not essential for baseline PTI outputs. Instead, they likely function as transcriptional hubs that bridge PTI and ETI, perhaps via enabling mutual potentiation of these two immune layers. This is consistent with our SETI-based transcriptomic profiling, which indicates that CBP60g and SARD1 control early immune gene reprogramming in an ETI-specific context.

One of the most striking features of the *gh* mutant is its exaggerated hypersensitive response (HR) upon ETI activation, despite reduced resistance. This phenotype sharply contrasts with most previously characterized HR regulators, which often show constitutive defense activation, pleiotropic developmental defects, or spontaneous lesion formation. Unlike *dnd1*, *hlm1*, or *lsd1*, *gh* plants grow normally and lack constitutive immunity, making them a valuable genetic model to uncouple cell death from resistance. Interestingly, while *eds5* mutants (also SA-deficient) exhibit strong susceptibility to the necrotrophic pathogen Botrytis cinerea, *gh* plants do not, despite comparable reductions in SA accumulation and enhanced HR. This suggests that HR in *gh* is qualitatively distinct from that in other SA-deficient mutants and may not be exploited by necrotrophs.

Transcriptomic profiling revealed that CBP60g and SARD1 exert both positive and negative transcriptional control. Clusters 5 and 8, strongly induced in Col-0 but attenuated in *gh*, included genes required for SA and NHP biosynthesis (*ICS1, PBS3, ALD1, FMO1, SARD4*), the *EDS1/PAD4* module, and numerous defence-related transcription factors, corresponding closely to known CBP60g and SARD1 binding targets. By contrast, clusters 6 and 10 were upregulated in *gh* and enriched for genes involved in Ca²⁺ signalling, cellular organization, and cell death execution, indicating that CBP60g and SARD1 normally constrain HR-promoting programs. Cluster 9, which was minimally affected in *gh* and showed little overlap with SARD1 ChIP targets, was enriched for WRKY, MYB, ANAC, ERF, bZIP, and bHLH motifs, suggesting CBP60g- and SARD1-independent transcriptional programs that sustain residual ETI. Together, these findings support a model in which CBP60g and SARD1 function as dual regulators, promoting defence gene activation while simultaneously restraining runaway cell death, such that their absence both weakens essential defence modules and releases repression on HR-associated pathways, producing the paradoxical phenotype of reduced resistance but enhanced HR.

We further tested this model by examining Nudix hydrolases, a family of nucleotide-metabolizing enzymes implicated in immune homeostasis. Among them, *NUDT7* was strongly induced upon ETI in Col-0 but showed markedly reduced expression in *gh*. Overexpression of *NUDT7* suppressed HR in both WT and *gh* backgrounds, but could not fully restore homeostasis in *gh*, demonstrating that loss of CBP60g and SARD1 disrupts multiple layers of negative regulation. This highlights that immune homeostasis depends on a combinatorial network of repressors, rather than on individual factors alone.

Our results broaden the view of CBP60g and SARD1 from SA-dependent activators to network-level integrators that balance immune activation and suppression. The exaggerated HR yet reduced resistance of *gh* mutants reveal that resistance and cell death, long considered tightly coupled, can be genetically and mechanistically separated. Future studies combining high-resolution temporal profiling, network modelling, and metabolomic analyses will be critical to identify upstream signals and downstream executioners of this atypical HR. More broadly, the dual roles of CBP60g and SARD1 underscore how plants achieve robust pathogen defence while minimizing self-inflicted damage, a principle of immune regulation with both mechanistic and translational significance.

## Acknowledgments

HC and PD acknowledge European Research Council Starting Grant “R-ELEVATION” (grant agreement: 101039824). JDGJ was supported by the Gatsby Charitable Foundation (UK).

## Author contributions

PD conceptualized and oversaw the inception of the research project. The experimental work was collaboratively conducted by HC, H-JS, and BPMN. Data analysis and figure generation were performed by HC and PD. JDGJ was involved throughout the project, providing valuable discussions that significantly shaped the research. HC and PD wrote the initial manuscript draft. All co-authors contributed to subsequent revisions. The final manuscript was prepared by HC and PD and was approved for submission by all authors.

## Data availability

The RNA-seq data for this study have been deposited in the European Nucleotide Archive (ENA) at EMBL-EBI under EMBL-EBI: PRJEB34958.

---

**Figure S1.**
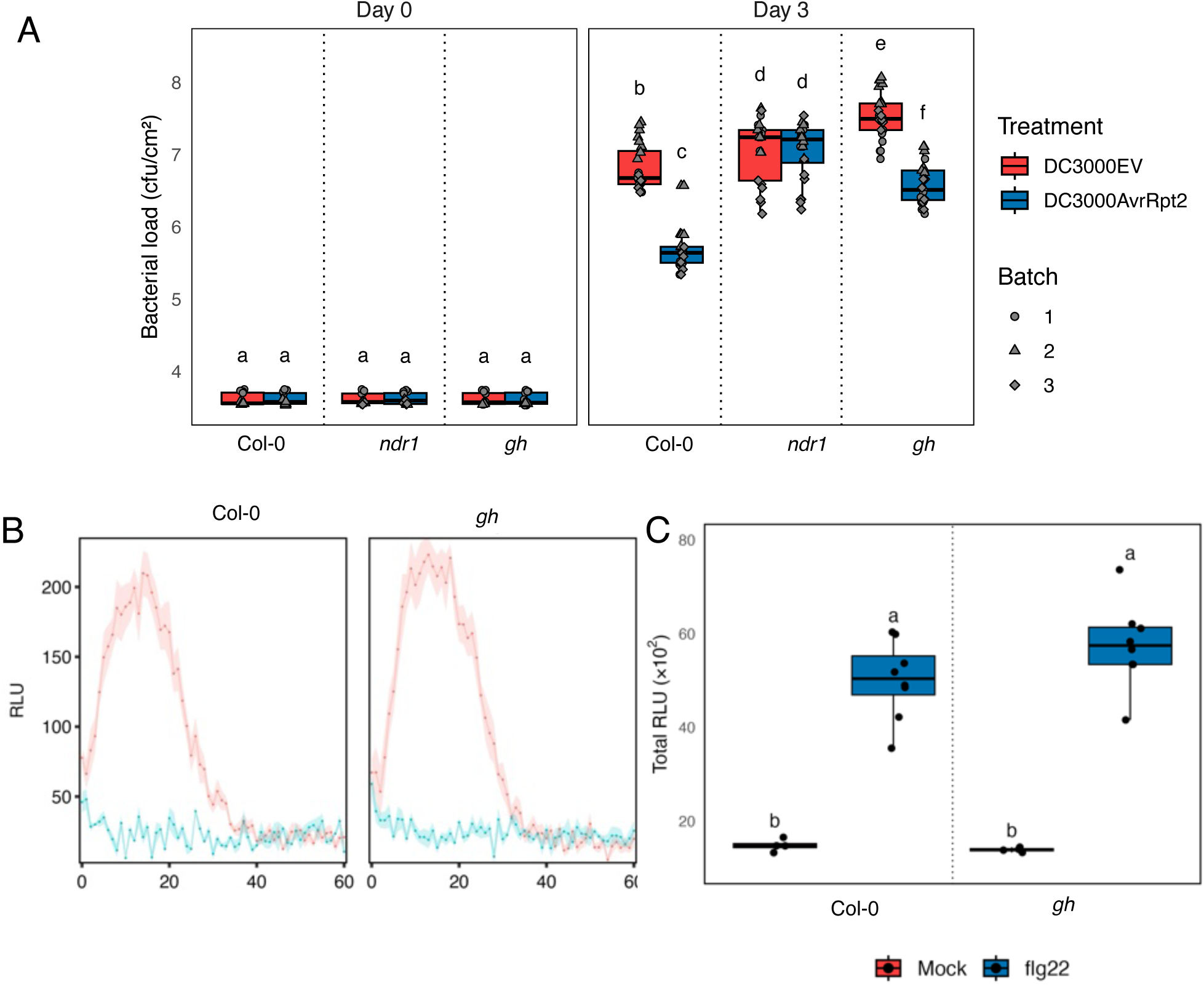
Contribution of CBP60g and SARD1 in PTI and ETI. **(A)** Leaves of Col-0 and the indicated mutant genotypes were infiltrated with *Pseudomonas syringae* pv. *tomato* DC3000 carrying either an empty vector (EV) or AvrRpt2. Bacterial growth was quantified as colony-forming units (CFU) at 0 and 3 days post-inoculation (dpi). Each data point represents two pooled leaves from a single plant. Black lines indicate mean values of technical replicates. Statistical differences are indicated by different letters (Tukey’s honestly significant difference [HSD] test, p ≤ 0.05). **(B)** Col-0 and *gh (cbp60g sard1)* were treated with mock or flg22 and ROS burst is measured. Statistical differences are denoted by different letters (Tukey’s HSD test, p ≤ 0.05). **(C)** Total RLU counts in Col-0 and *gh (cbp60g sard1)* plants. Statistical differences are denoted by different letters (Tukey’s HSD test, p ≤ 0.05).

**Figure S2.**
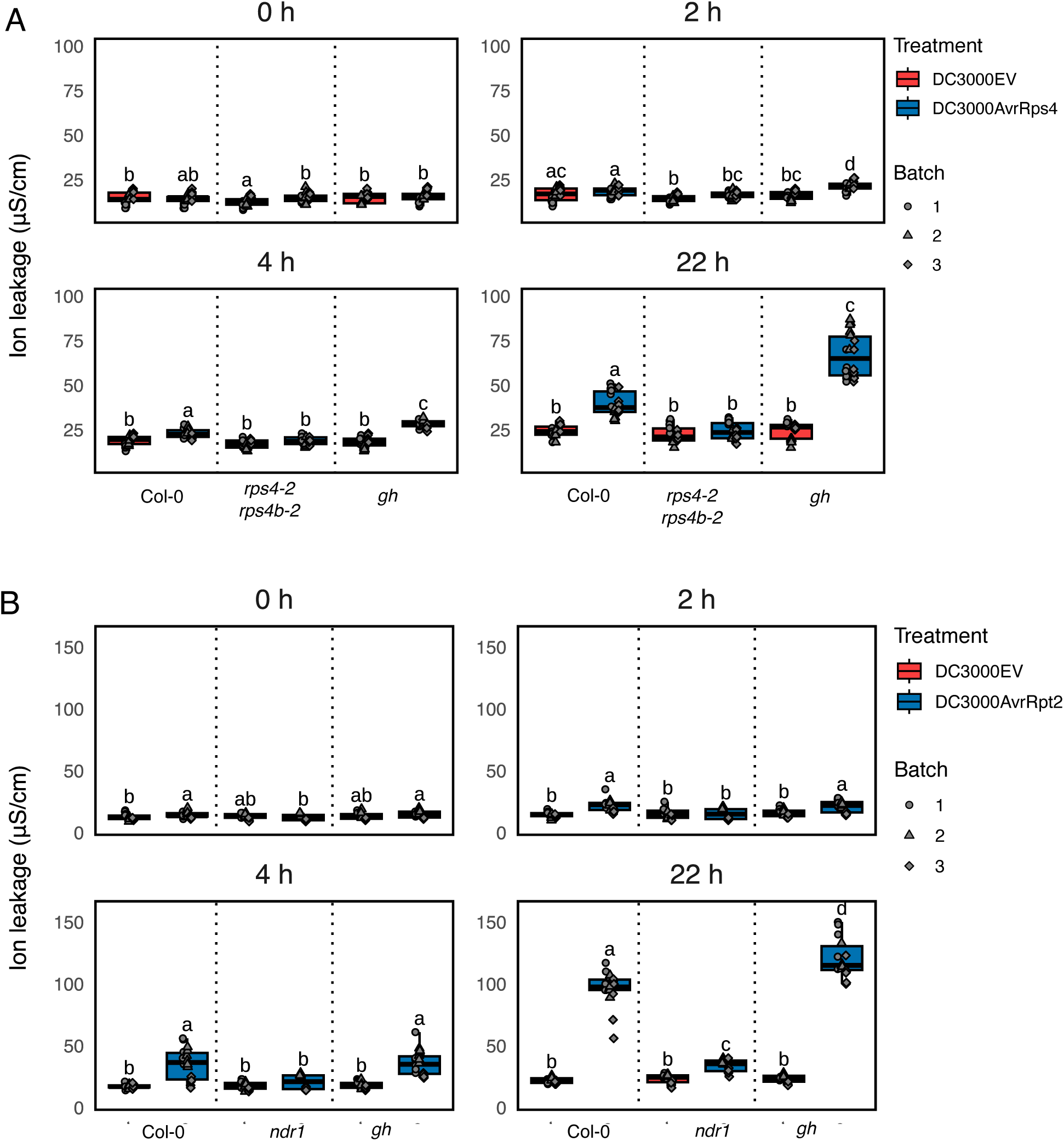
Loss of CBP60g and SARD1 enhances ETI^AvrRps4^-induced HR. **(A) and (B)** Ion leakage in indicated mutants infiltrated with mock, E2, or *Pseudomonas syringae* DC3000 (empty vector or AvrRps4) and (empty vector or AvrRpt2) measured at 0–22 h post-infiltration. Different letters indicate statistical significance (Tukey’s HSD test, p ≤ 0.05).

**Figure S3.**
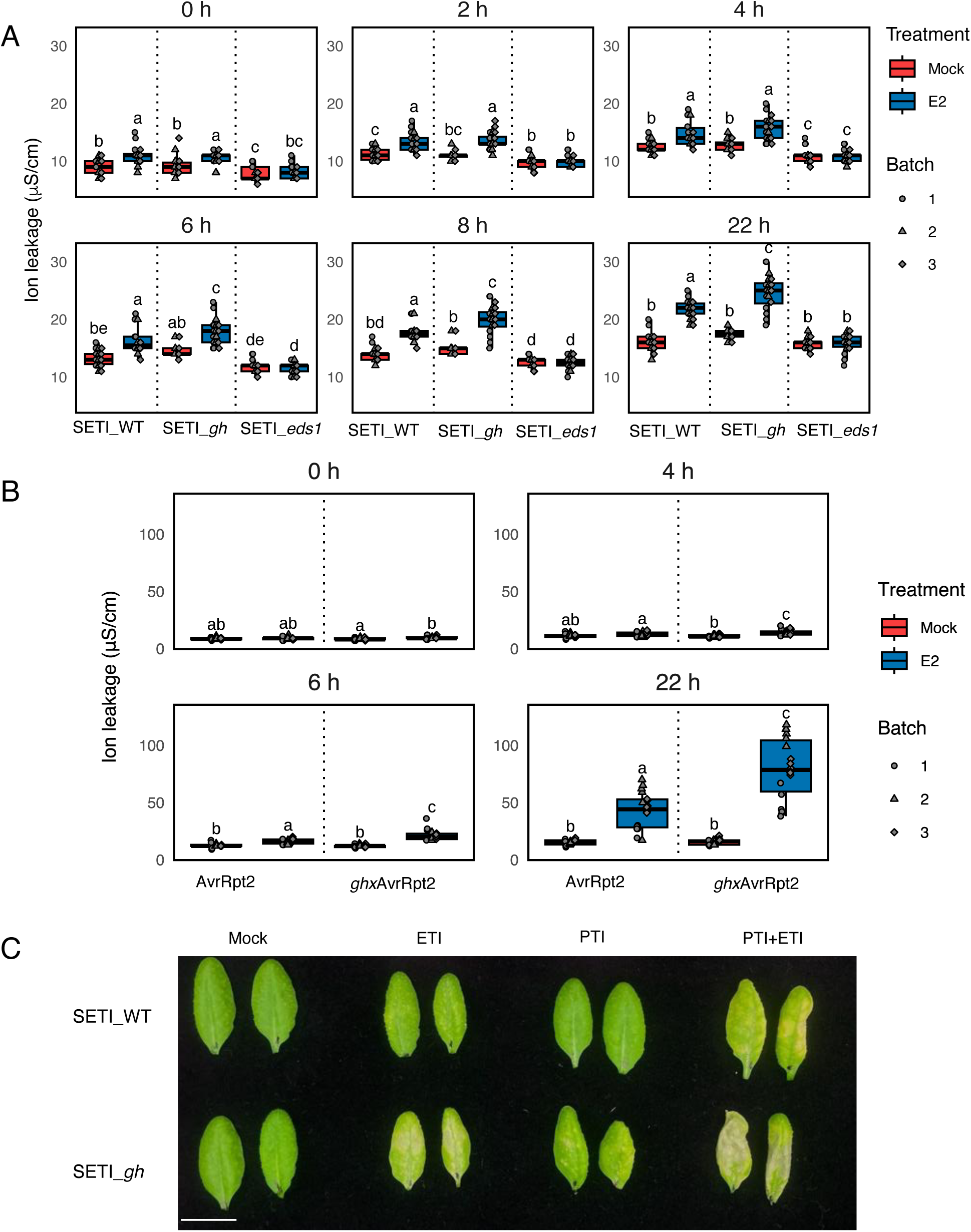
Loss of CBP60g and SARD1 enhances ETI-induced hypersensitive response (HR) **(A)** and **(B)** Ion leakage measurements from the indicated mutants infiltrated with mock or E2 at the specified time points. Different letters indicate statistically significant differences (Tukey’s HSD test, *p* ≤ 0.05). **(B)** HR phenotypes of SETI_WT and SETI_*gh* (SETI_*cbp60g sard1*) plants infiltrated with mock, E2 (ETI), DC3000 hrcC⁻ (PTI), or PTI+ETI treatments, photographed 3 days post-infiltration. Scale bar = 1 cm

**Figure S4.**
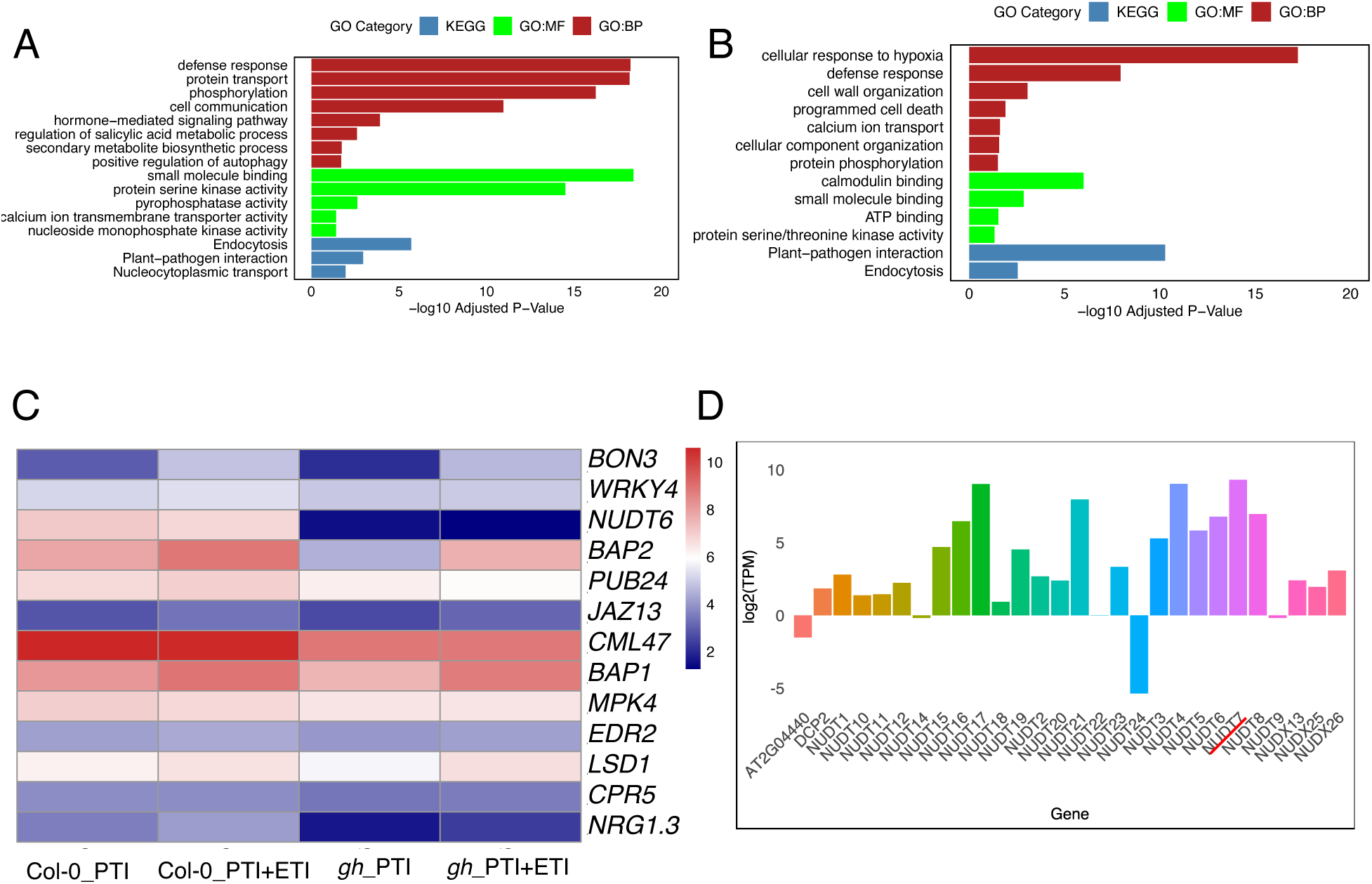
Negative regulation of immune components by CBP60g and SARD1. **(A, B)** GO enrichment analysis of clusters 6 and 10 using g:Profiler, categorized into biological process (GO:BP), cellular component (GO:KEGG), and molecular function (GO:MF) terms (FDR ≤ 0.05, Benjamini–Hochberg correction). **(C)** Heatmap showing expression of representative negative immune regulators in WT and *gh* (*cbp60gsard1*). **(D)** Expression profiles of NUDT family members following PTI+ETI treatment. Nudix7 is highlighted with red line.

**Figure S5.**
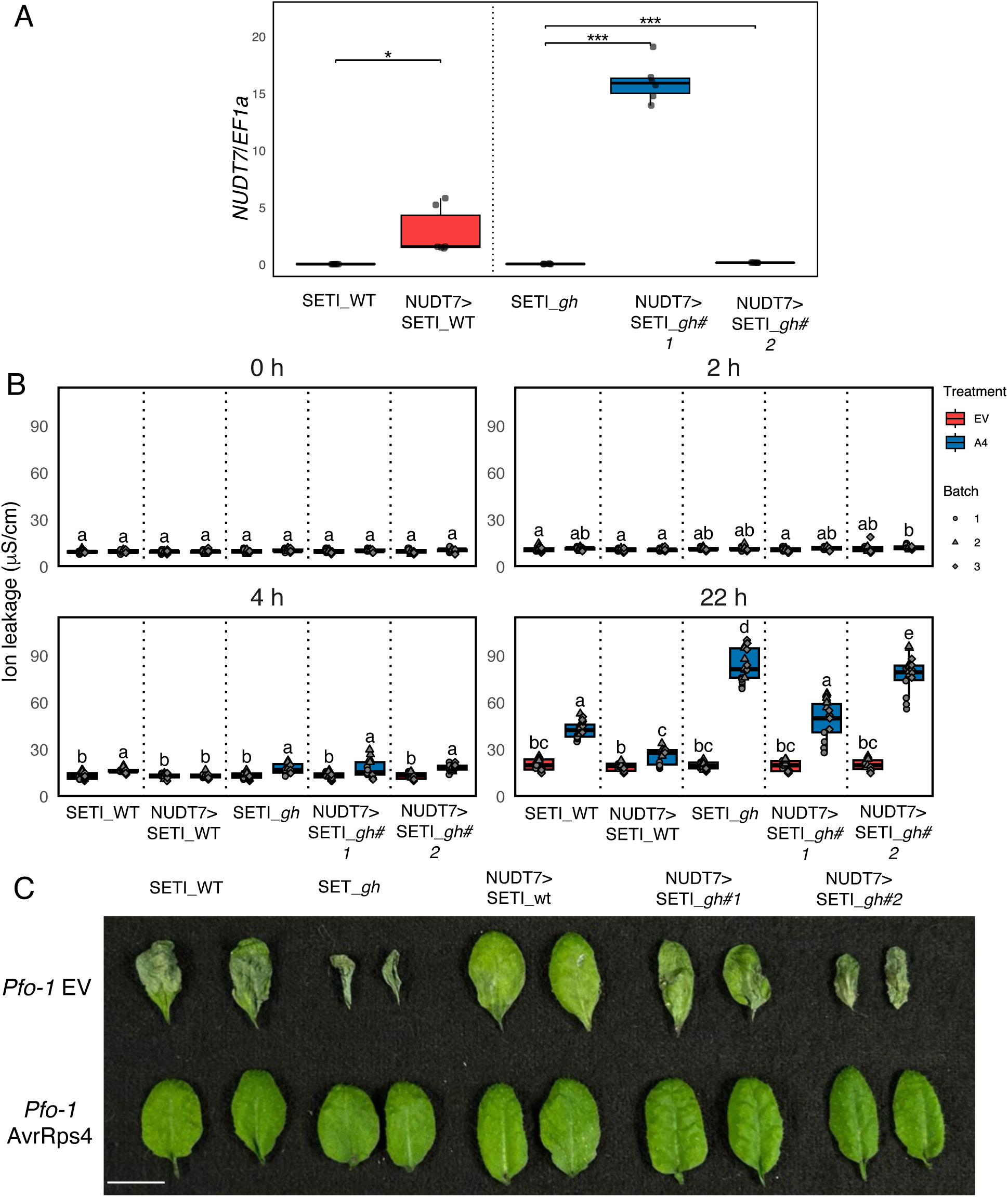
NUDT7 overexpression cannot fully rescue enhanced HR in. (A) Leaves from Five-week-old plants of SETI_WT, SETI_*gh* (SETI_*cbp60g sard1*) along with the overexpression lines of NUDT7 in SETI_WT and SETI_*gh* background were harvested for RNA extraction and qPCR. *NUDT7* expression levels are shown relative to *EF1α.* Scale bar = 1 cm. Asterisk indicate significant difference (t-test, p ≤ 0.05). (#1 and #2 independent lines). (B) Ion leakage in indicated mutants infiltrated with *Pseudomonas syringae* DC3000 (empty vector or AvrRps4) measured at 0–22 h post-infiltration. Different letters indicate statistical significance (Tukey’s HSD test, p ≤ 0.05). (C) HR phenotypes of indicated genotypes infiltrated with *Pf0-1* carrying empty vector or AvrRps4 at 3 days post-infiltration.

